# Assessment of the suitability of marker fragments DNA sequences of the internal transcribed spacer region (ITS), beta-tubulin gene (TUB) and elongation factor (EF1) genes for diagnostics of *Fusarium tricinctum* species complex in Russian in grain products

**DOI:** 10.1101/2021.02.15.431223

**Authors:** A. Shchukovskaya

## Abstract

By results of laboratory researches from plant samples of grain crops species *Fusarium avenaceum, Fusarium tricinctum* and *Fusarium acuminatum* related to *Fusarium tricinctum* species complex were isolated and identified, based on cultural and morphological characteristics, molecular-genetic analysis of 3 DNA fragments was performed: internal transcribed spacer (ITS) site, beta-tubulin gene site (TUB), elongation factor gene site (EF1) showed insignificant intraspecific and interspecific variability at ITS and EF1 sites, which may negatively affect the reliability of target species identification, TUB site was more effective for distinguishing and identifying the studied species, due to significant intraspecific and interspecific variability

## INTRODUCTION

The export of crop products is becoming increasingly important in the economy of the Russian Federation. In addition, countries importing Russian grain impose certain phytosanitary requirements that must be met by products supplied to the buyer country.

The most significant phytopathogens of grain crops are micromycetes of the genus Fusarium, which can cause damage to all vegetative organs and contaminate grain with mycotoxins during vegetation and storage [2]. The species *F. acuminatum* Ellis & Everh., *F. avenaceum* (Fr.) Sacc., Sylloge, *F. tricinctum* (Corda) Sacc. affecting ears, cobs, grain products, causing root rot and being a part of *Fusarium tricinctum* species complex are subject to phytosanitary requirements in several countries where russian grain products are exported.

Identification of these species by classical mycological methods is not always reliable; therefore, molecular diagnostic methods, namely the selection and evaluation of the efficiency of marker DNA sequences for the identification of analyzed species, are of particular interest [2].

### Material and methods

We analyzed pure cultures of *F. avenaceum*, *F. tricinctum*, and *F. acuminatum* isolated from plant samples of grain crops (ears, grains, leaves, and stems) infected with Fusarium (Fig. 1).

**Figure 1.**
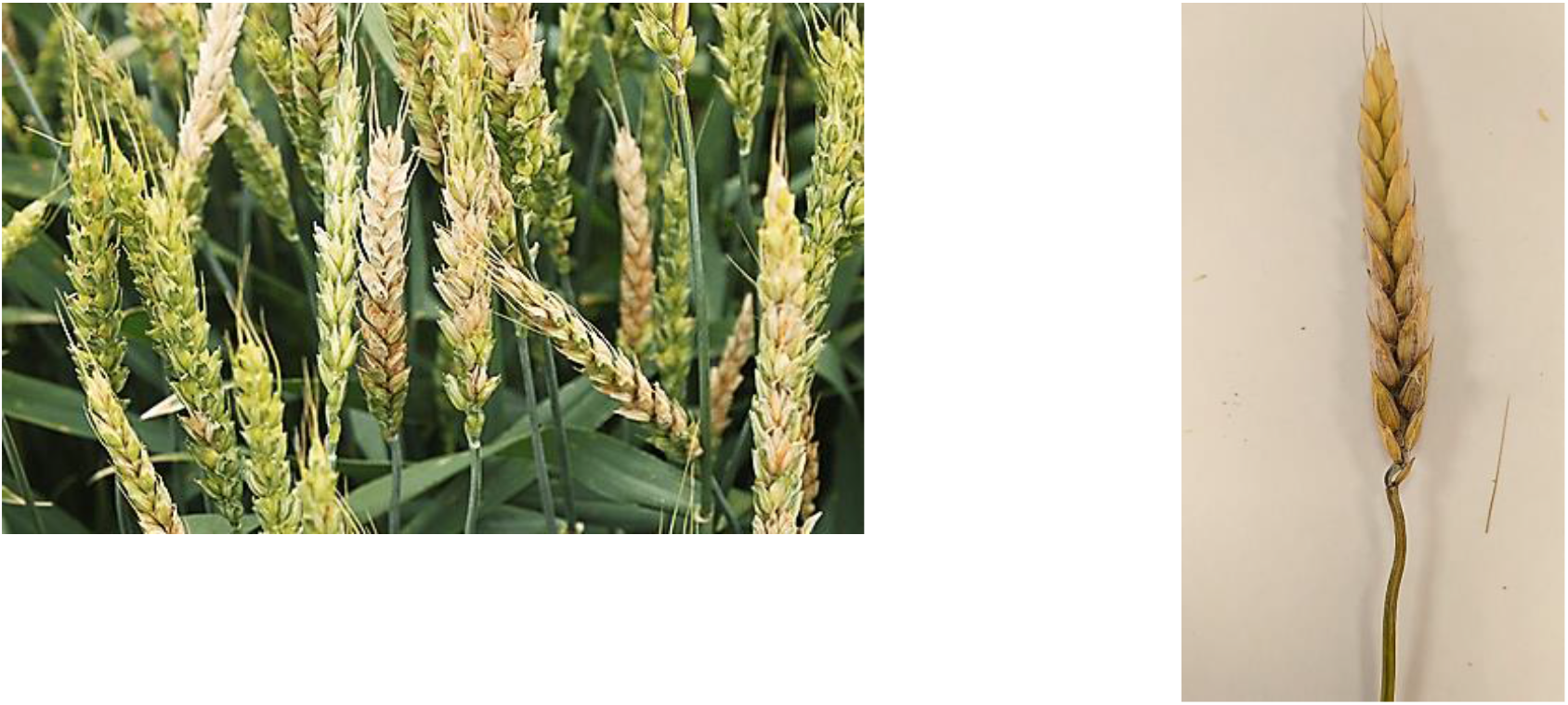
Ears of winter wheat infected with Fusarium [author’s picture].

Further identification of the target species was performed using a molecular genetic method. Pathogen identification was based on determining the nucleotide sequences of the transcribed spacer sites, the beta tubulin gene, and the elongation factor gene. Amplification of these loci was carried out with the primers listed in Table 1.

**Table 1.**
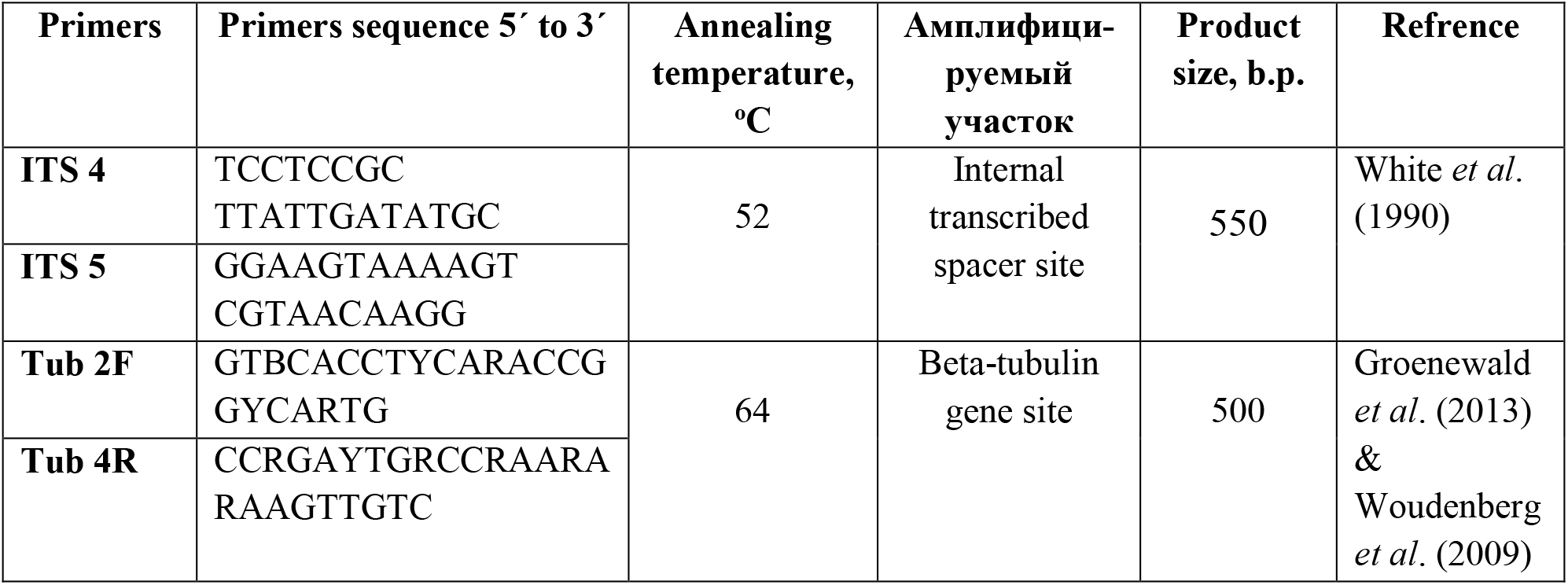
List of primers used in the work.

Nucleotide sequences were determined from the obtained PCR products by Sanger Sequencing on on an Applied Biosystems 3500 device (Thermo Scientific, USA). Nucleotide sequences were aligned using the Clustal W algorithm in BioEdit software. Comparative analysis of the sequenced nucleotide sequences with those deposited in GenBank was performed using the BLAST program on the NCBI website [6]. Dendrograms were constructed using the MEGA-X program (MEGA-X: Molecular Evolutionary Genetics Analysis version 10.0.5 (Tamura, Stecher, and Kumar, 1993-2020).

Original sequences of target species and nucleotide sequences (target and other species of the genus *Fusarium*) from the NCBI database were used: original nucleotide sequences of DNA sites (ITS, TUB) - *F. avenaceum* (F6t), *F. acuminatum* (F87), *F. tricinctum* (F11). Nucleotide sequences of the site (EF1) of the target and other species of the genus *Fusarium* are from the NCBI database.

## RESULTS AND DISCUSSION

The sequence similarity dendrogram for the internal transcribed spacer region shows five distinct clusters with high bootstrap test support (98-99%). Species of *F.solani, F.sporotrichioides, F.oxysporum, F.graminearum*, and species of *Fusarium tricinctum* complex formed separate groups (Figure 2). The most heterogeneous cluster is represented by species of the *Fusarium tricinctum* complex, which are characterized by intraspecific differences, making it impossible to identify them. This site is not effective for distinguishing and determining target species

**Figure 2.**
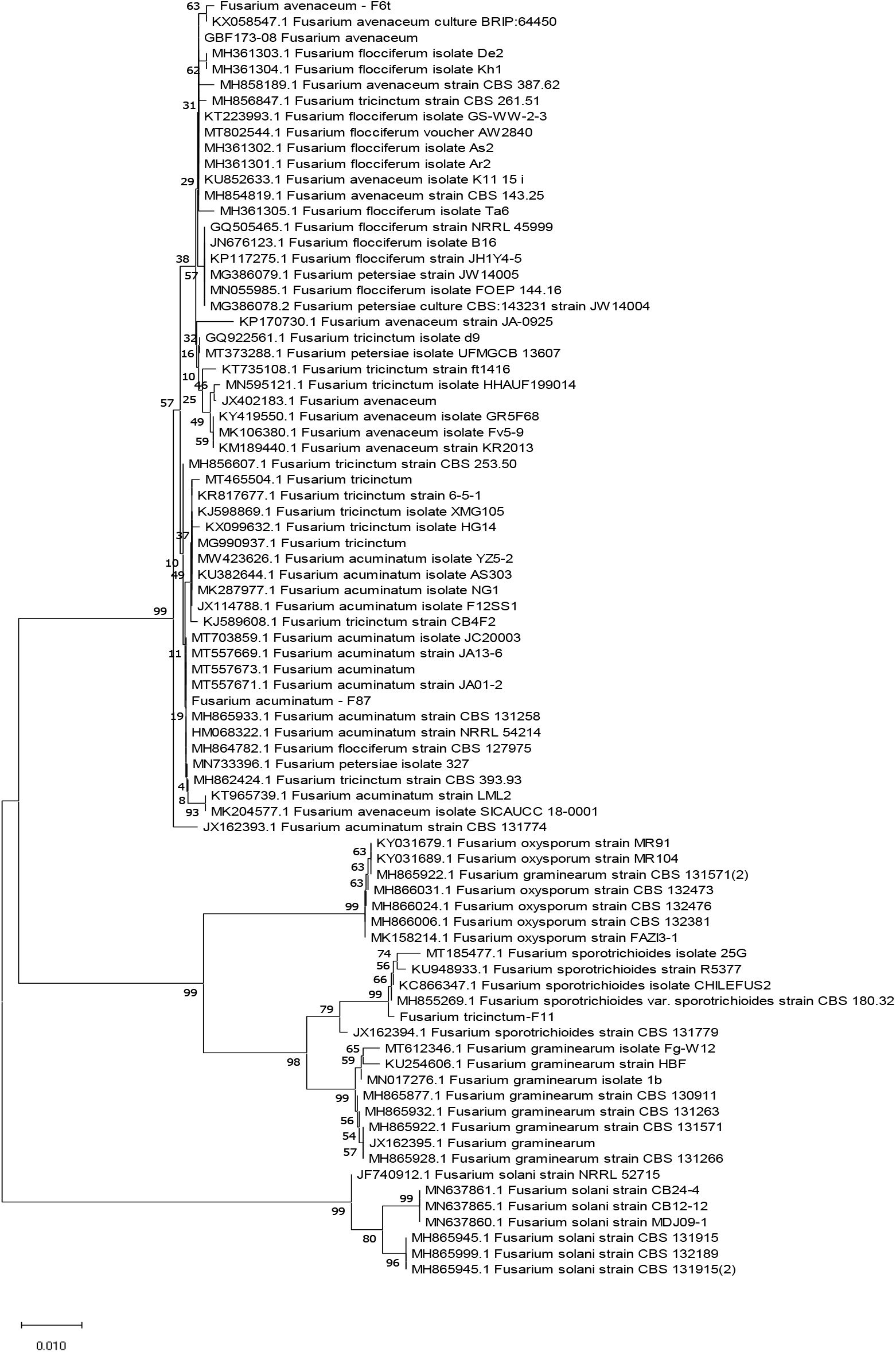
Dendrogram of similarity and difference of sequences of the internal transcribed spacer site of the studied *Fusarium* species (species name and sample number are given), constructed using the nearest neighbor (NJ) linking method. Stability determined by bootstrap analysis (1000 alternative dendrograms), given as a percentage above the branches

The sequence similarity dendrogram for the beta-tubulin gene fragment also shows five distinct clusters with high support of the bootsrap test (76%-100%). The division of target species within the complex is clearer: isolates of *F. acuminatum* species represent a separate group; species of *F. avenaceum* and *F. tricinctum* are also separated (Figure 3). The beta-tubulin gene site is more effective in distinguishing and identifying the species under study.

**Figure 3.**
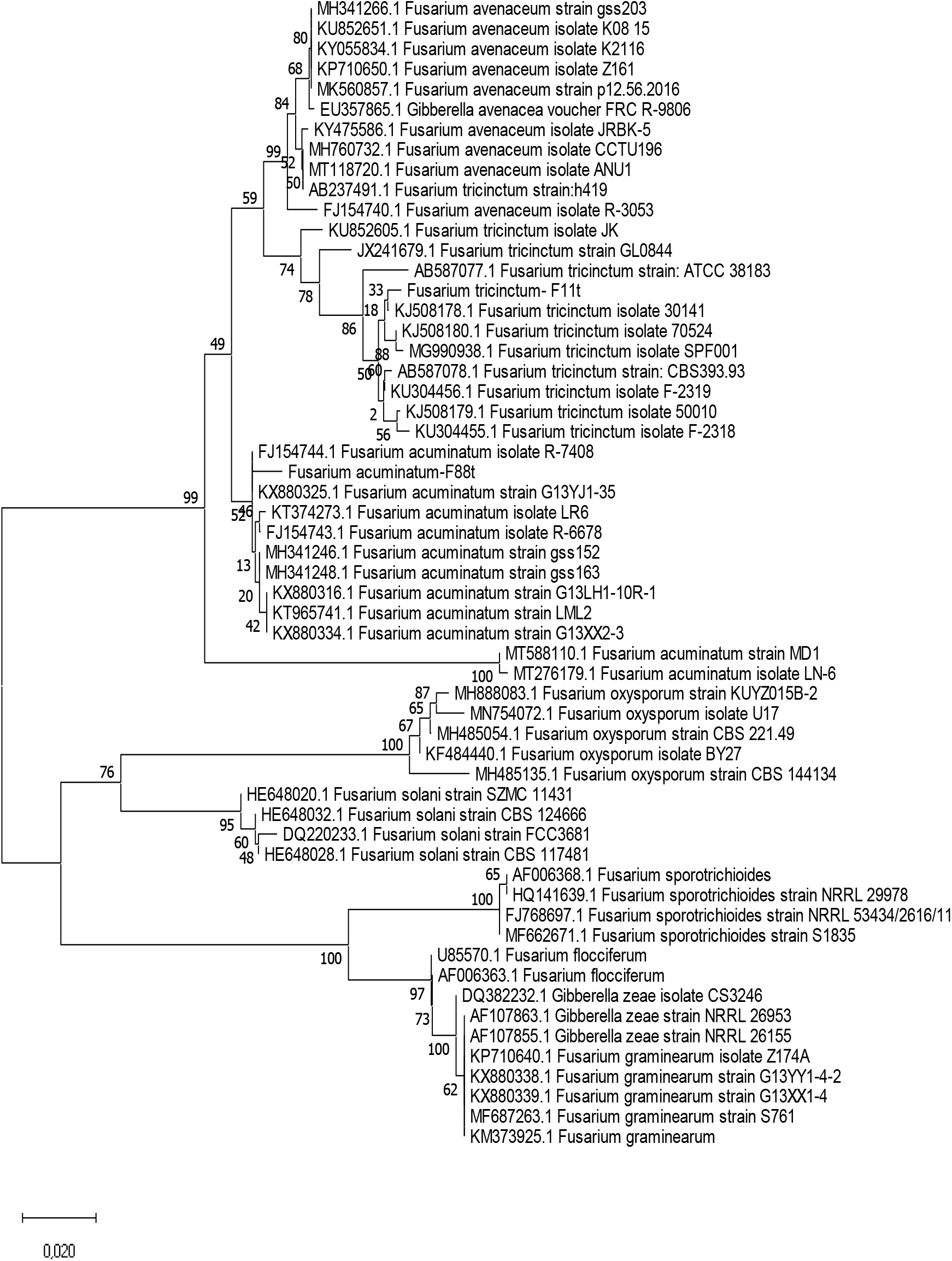
Dendrogram of similarity and difference of TUB gene fragment sequences of the studied Fusarium species (species name and sample number are given) constructed using the nearest neighbor (NJ) linking method. Stability determined by bootstrap analysis (1000 alternative dendrograms), given as a percentage above the branches * * The sequences of the beta-tubulin gene region for *F. petersiae* are not shown in the dendrogram because they are not available in the GenBank database.

The sequence similarity dendrogram for the elongation factor gene fragment also shows 5 separate clusters with a bootstrap test support of 63%-100% (Figure 4). A clear separation of the target species and other species of the genus *Fusarium: F. solani, F. oxysporum, F. graminearum*, and *F. sporotrichioides* occurs according to this gene region. However, the analyzed species are less separated from each other; for example, species of *F. flocciferum* and *F. petersiae* are not distinguished from each other and are combined into one cluster, which in turn can negatively affect the reliability of target species determination. The elongation factor gene site can be used to distinguish and identify target species, but not for all species.

**Figure 4.**
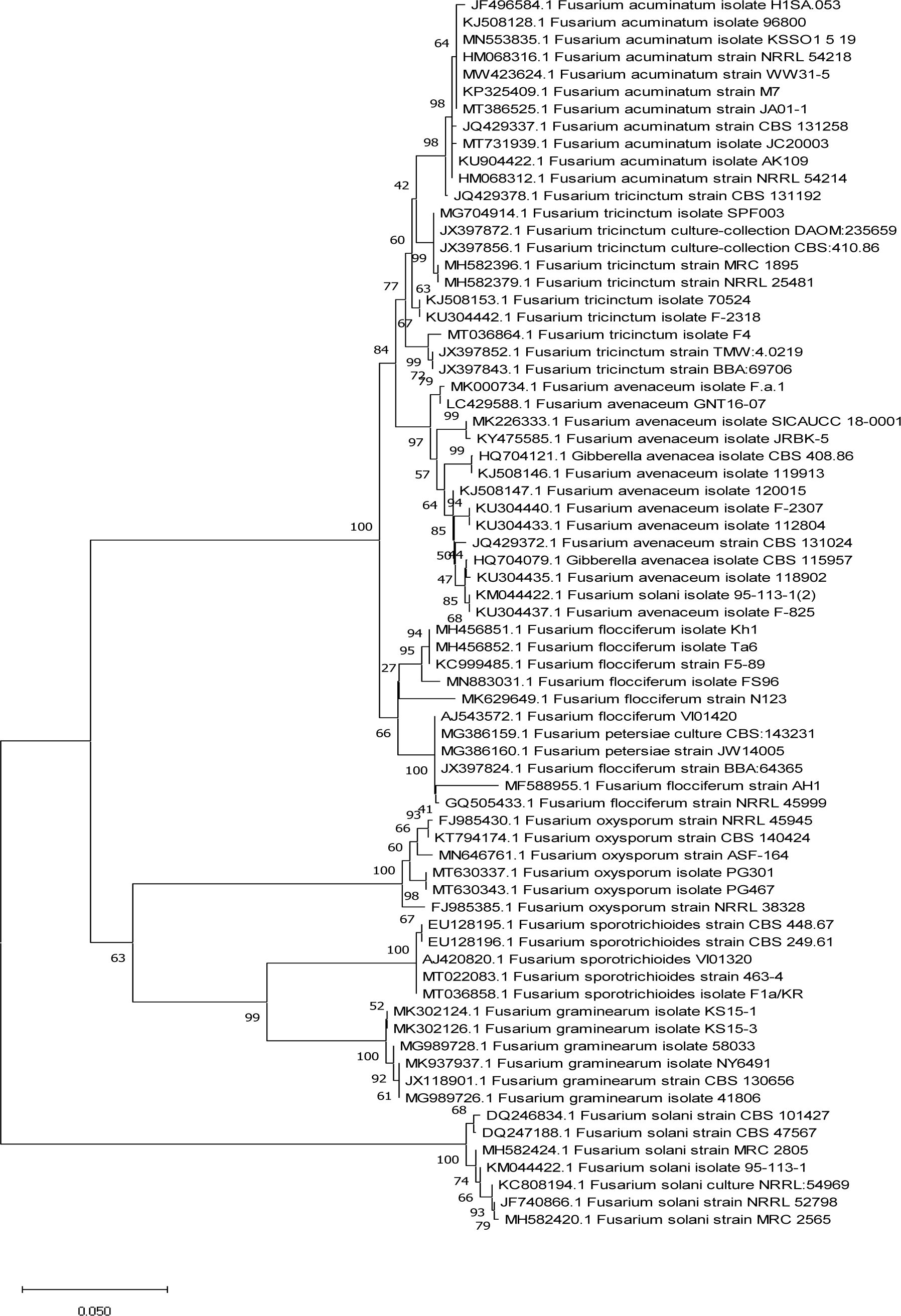
Dendrogram of similarity and difference of EF1 gene fragment sequences of the studied *Fusarium* species (species name and sample number are given) constructed using the nearest neighbor (NJ) linking method. Stability determined by bootstrap analysis (1000 alternative dendrograms) is given as a percentage over the branches

In this regard, multilocus analysis and the search for efficient marker sequences is a relevant direction.

